# Predictability of salient distractor increases top-down control in healthy younger and older adults

**DOI:** 10.1101/617712

**Authors:** Marleen Haupt, Natan Napiórkowski, Christian Sorg, Hermann J. Müller, Kathrin Finke

**Affiliations:** General and Experimental Psychology, Department of Psychology, Ludwig-Maximilians-Universität München, Germany; Graduate School of Systemic Neurosciences (GSN), Ludwig-Maximilians-Universität München, Germany; Hans-Berger Department of Neurology, University Hospital Jena, Germany; Department of Neuroradiology, Klinikum rechts der Isar, Technische Universität München, Germany; Department of Psychiatry and Psychotherapy, Klinikum rechts der Isar, Technische Universität München, Germany

**Author notes:** Corresponding author Marleen Haupt, Department of General and Experimental Psychology, Ludwig-Maximilians-Universität, Leopoldstrasse 13, 80802 München, Germany.

**Keywords:** Healthy Ageing, Top-down Control, Distractor Predictability, Visual Attention, Active Perception

## Abstract

Younger adults are able to shield attentional selection against distractors when they have pre-knowledge about the upcoming distractor location. For older adults, who suffer from an overall decrease in attentional capacity and who are, in addition, particularly prone to attentional capture, such an adaptive shielding ability would be of particular importance. However, it is an open question whether healthy older adults can utilise the predictability of distractor locations to improve top-down controlled selection to the same degree as younger adults. The theory of visual attention (TVA) framework provides a systematic way to measure an individual’s efficiency of top-down control. The present study combined a TVA-based partial-report paradigm with abrupt-onset cues rendering the indicated location highly salient in a bottom-up fashion. Experiment 1, in which (on cued trials) the cue was invariably followed by a distractor at the cued location, showed that the cueing increased the weight of the distractor in the competition for selection compared to uncued distractors (on trials without a cue). In Experiment 2, the probability with which the abrupt-onset cue indicated the upcoming distractor location (1/3 vs. 2/3 of trials) was manipulated between experimental blocks. Participants were able to learn these statistical contingencies and exert top-down control more efficiently in blocks with highly valid distractor location cues, as compared to low-validity blocks. This finding suggests that, even though abrupt-onset spatial cues increase the attentional weights of distractors, participants can acquire and use pre-knowledge about the likelihood that a distractor will appear at an indicated location to down-weight the bottom-up attentional-capture signal. This ability turned out to be comparable across age groups, suggesting that efficient use of predictive information to shield against distracting information is preserved in normal ageing.

## Introduction

Our visual environment is too complex for us to process all objects that compete for our limited visual processing resources at any given time. Moreover, as the available resources decline over the lifespan, the amount of information that can actually be processed is even more reduced in older compared to younger adults (e.g. McAvinue et al., 2012; Ruiz-Rizzo et al., 2019; Salthouse, 1996). According to the biased-competition view of attentional selection (Desimone & Duncan, 1995), an interaction of stimulus-related bottom-up salience and observer-related top-down mechanisms lead to a biased (attentional) weighting of the objects in the visual array (Bundesen, 1990; Corbetta & Shulman, 2002; Desimone & Duncan, 1995; Theeuwes, 2010). An example of a bottom-up controlled effect is attentional capture by highly salient (but task-irrelevant) ‘distractors’ in the display, which hampers responding to the (task-relevant) target (e.g.,Theeuwes, 1991, 1992). However, younger adults can mitigate the detrimental effects of strong bottom-up signals by utilising prior information (‘expectancies’) about where an upcoming distractor is likely to occur. That is, they can improve top-down controlled weighting of stimuli, and thus reduce distractor interference, when they can make valid predictions about the (likely) location of an upcoming distractor based on spatial pre-cues (Awh, Matsukura, & Serences, 2003; Chao, 2010; Havlíček, Müller, & Wykowska, 2018; Noonan et al., 2016; Ruff & Driver, 2006; Watson & Humphreys, 1997) or based on statistical learning of the spatial distribution of distractors in within a search display (Goschy, Bakos, Müller, & Zehetleitner, 2014; Sauter, Liesefeld, & Müller, 2019; Sauter, Liesefeld, Zehetleitner, & Müller, 2018). For older adults who are affected by age-related reductions in attentional capacity, a preserved ability to use predictive information about where distracting stimuli might appear would be of particular importance. An adaptive increase in the efficiency of top-down control would (at least partially) compensate for the capacity limitations, as better shielding against distracting information would reduce the information overload. However, prior evidence on the use of predictive information in attentional selection by older adults is mixed. Older individuals have been reported to be more susceptible to attentional capture compared to younger adults, suggesting that their attentional selection processes are more controlled by bottom-up factors (Healey, Campbell, & Hasher, 2008; Kramer, Hahn, Irwin, & Theeuwes, 2000; Phillips & Takeda, 2010). But there are some studies suggesting that older participants remain, nevertheless, able to efficiently use spatial information about (likely) target locations in search displays to reduce attentional capture (Whiting, Madden, & Babcock, 2007), improve search performance (Madden et al., 2014; Madden, Whiting, Cabeza, & Huettel, 2004), and decrease susceptibility to noise (Whiting, Sample, & Hagan, 2014). To our knowledge, however, it has not yet been systematically addressed whether the ability to use predictive information about the location of distractors is preserved, too, in healthy older adults.

The theory of visual attention (TVA; Bundesen, 1990) provides a systematic framework for quantifying an individual’s efficiency of top-down control. Applying this framework (and attendant paradigms), the present study investigated whether top-down control is actively modulated according to observers’ expectancies about where upcoming distractors might appear. Specifically, we assessed whether observers would be able to reduce distractor interference in conditions in which the probability that a distractor is presented at a salient location is high versus low, and whether the ability to modulate top-down control is influenced by ageing.

TVA assumes that visual categorization allows for the simultaneous selection and recognition of stimuli (Bundesen, 1990): once the decision that an object belongs to a certain category is made, the object is encoded into a capacity-limited visual short-term memory (vSTM) store, making it available for conscious report. However, with multiple objects in the visual array, there is a competitive race for access to and representation in vSTM: only those objects that enter before the store is filled to capacity will be represented and available for report. The probability for a given object to be represented in vSTM depends on its relative attentional weight (w). Some objects receive higher weights and are, thus, favoured for selection, based either on automatic, bottom-up, or on intentional, top-down, factors. Estimates of individuals’ efficiency of top-down control are derived by modelling verbal report accuracy in a psychophysical partial-report task of briefly presented letter arrays containing both target and distractor items distinguished by different colours. TVA-based modelling of report accuracy across different (stimulus) conditions yields separate estimates of the attentional weights for target letters (to be encoded into vSTM and reported) and distractor letters (not to be encoded) (Bundesen, 1990; Duncan et al., 1999). TVA parameter α – that is, the ratio of distractor weights to target weights for a given individual – provides an index of this individual’s efficiency of top-down control.

The present study combined such a TVA-based partial-report paradigm with a distractor-related spatial-cueing procedure. The cues were abrupt-onset stimuli (a bright square around a position within the (subsequent) letter array) presented shortly before the appearance of a distractor (or target) letter at the respective location, thus rendering the respective location highly salient in a bottom-up fashion. Our analyses focused on trials in which the cue was followed by a distractor, in order to compare the ability to prepare for the distractor and shield against interference based on the likelihood of a distractor appearing at the location made salient by the cue. The probability with which the distractor appeared at the cued location was varied block-wise: 1/3 (of trials) in low-probability blocks, and 2/3 in high-probability blocks. Importantly, the bottom-up salience of the distractors (i.e., the boost they received by the preceding abrupt-onset cue at their location) was equal in both conditions. Accordingly, comparing estimates of top-down control (parameter α) allowed us to assess whether observers would be able to improve top-down control when the cue indicated the location of a distractor with a high probability, as compared to a low probability. The bottom-up saliency boost produced by the cue would result in equal (bottom-up) attentional weights of the distractors in both conditions, and, thus, top-down control parameter estimates based on distractor and target weights would be expected to be equal, too. However, if observers can make efficient use of the probabilistic information conveyed by the cue (i.e., about the likelihood of a distractor appearing at the cued location), this should lead to a (top-down) down-modulation of distractor weights and, thus, better top-down control in the high-probability versus the low-probability condition. As the TVA-based partial report allows for a parametrization of top-down control based on non-speeded verbal report accuracy, this assessment is especially appropriate for healthy older adults, who typically exhibit an overall slowing of their motor performance (Shalev, Humphreys, & Demeyere, 2016).

In sum, the present study set out to investigate whether younger and older (healthy) participants can utilise pre-knowledge about the location of upcoming distractors to effectively adjust their top-down control settings and thus reduce distractor interference.

## Exneriment 1

Our study aim was to examine whether observers can make use of the probabilistic information conveyed by abrupt-onset cues to (top-down) reduce the weight of distractors at this location (in the competition for selection into vSTM). For this, we first had to establish the (bottom-up) effect of the cues on the distractor weights in a preliminary experiment (Experiment 1). Given that abrupt-onset cues render the (distractor) stimulus that occurs there more (bottom-up) salient compared to a (distractor) stimulus at the same location not ‘highlighted’ by a cue, the attentional weight for the distractor should be higher relative to that for the target (which never occurs at the location of the cue) when the distractors occurs at the cued location versus when it occurs at the same location in the absence of a cue. As a result, observers should be less able to ignore a distractor in the cued condition (because it gains a higher weight by virtue of being made salient by the cue), which would be expressed in a higher value of the TVA top-down control parameter α (i.e., the ratio of the distractor to the target weights). Experiment 1 was designed to test this prediction using an abrupt-onset cue in one condition (in which a distractor was in competition with a target), which was 100% predictive (or ‘valid’) with respect to the location of the distractor. Establishing this effect was a requirement for the main experiment, Experiment 2, which went on to introduce a cue-validity manipulation to examine for a top-down modulation of the distractor weight at the cued location.

## Method

### Participants

Eight healthy, right-handed volunteers (2 females) took part in the pilot study; their mean age was 25.9 (SD=1.6) years, and they had 12 years (SD=0.5) of education, on average. They all reported normal (or corrected-to-normal) vision, including normal colour vision. The study was approved by the Ethics Committee of the Department of Psychology at Ludwig-Maximilians-Universität (LMU) München.

### TVA-based partial-report paradigm

All participants underwent a TVA-based partial-report paradigm adopted from Finke et al. (2005), with an additional cueing manipulation (see below). On each trial, they were presented with a display of four (equidistant) gray squares (2.20° of visual angle), one in each display quadrant, located 5.72° of visual angle from a central fixation marker (a white circle, 0.79° of visual angle in diameter, with a dot inside; see Figure 1). The squares served as placeholders, indicating the four possible locations at which stimuli could appear. Three different types of stimulus displays (trials) were used in the partial-report task: trials containing (i) a single target letter, (ii) two target letters, and (iii) one target plus one distractor letter. In displays containing two stimuli, these were arranged either horizontally (one on each side) or vertically (both on the same side), but never diagonally. This resulted in 16 different display configurations (see Figure 1B). Targets were defined by being red and distractors by being blue, with luminance equated between the two colours (CIE_red_ = [0.49, 0.515, 0.322], CIE_blue_ = [0.49, 0.148, 0.068]); the screen background was black. All letter stimuli were 1.06° of visual angle heigh and 0.88° wide. Participants were instructed to verbally report the target letters they had recognized, without stress on response speed or report order (if they had recognized multiple targets). At the end of each completed test block, participants were presented with a scale illustrating whether their report accuracy ranged between 70% and 90% correct, the aimed-for level of accuracy. If their scores were higher or lower, they were provided with adapted instructions for the next block: if accuracy was above 90%, they were told by the experimenter to attempt to report more letters, even if they were not absolutely confident that their report is correct; if accuracy dropped below 70%, they were told to try to refrain from guessing and report only letters they had recognized with a high degree of confidence. After entering all reported letters on the computer keyboard, the experimenter started the next trial by pressing the space bar of the computer keyboard. Each trial consisted of one or two letters randomly drawn, without replacement, consisting of the following set of 23 letters {ABCDEFGHJKLMNOPRSTUVWXZ} (so no letter could appear twice in a display). All letters were presented for a brief, individually determined exposure duration and were then immediately masked with squares (1.72° of visual angle) filled randomly with red and blue miniblobs (CIE_red_ = [4.86, 0.605, 0.318], CIE_blue_ = [2.99, 0.162, 0.088]), which disappeared after 500ms.

**Figure 1.**
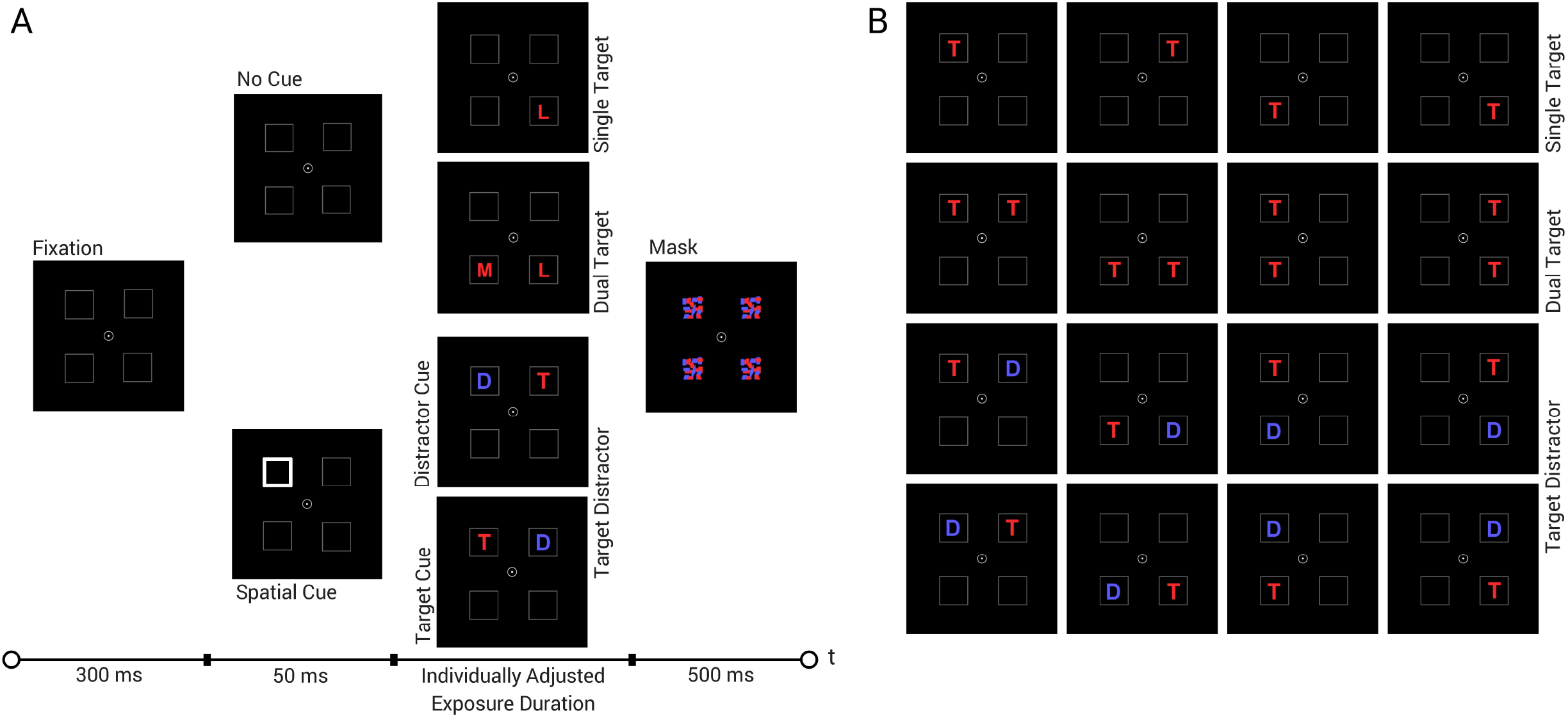
Partial Report Task: A) Trial sequence with examplary letters; B) 16 possible display configurations of targets (T) and distractors (D).

**Figure 2.**
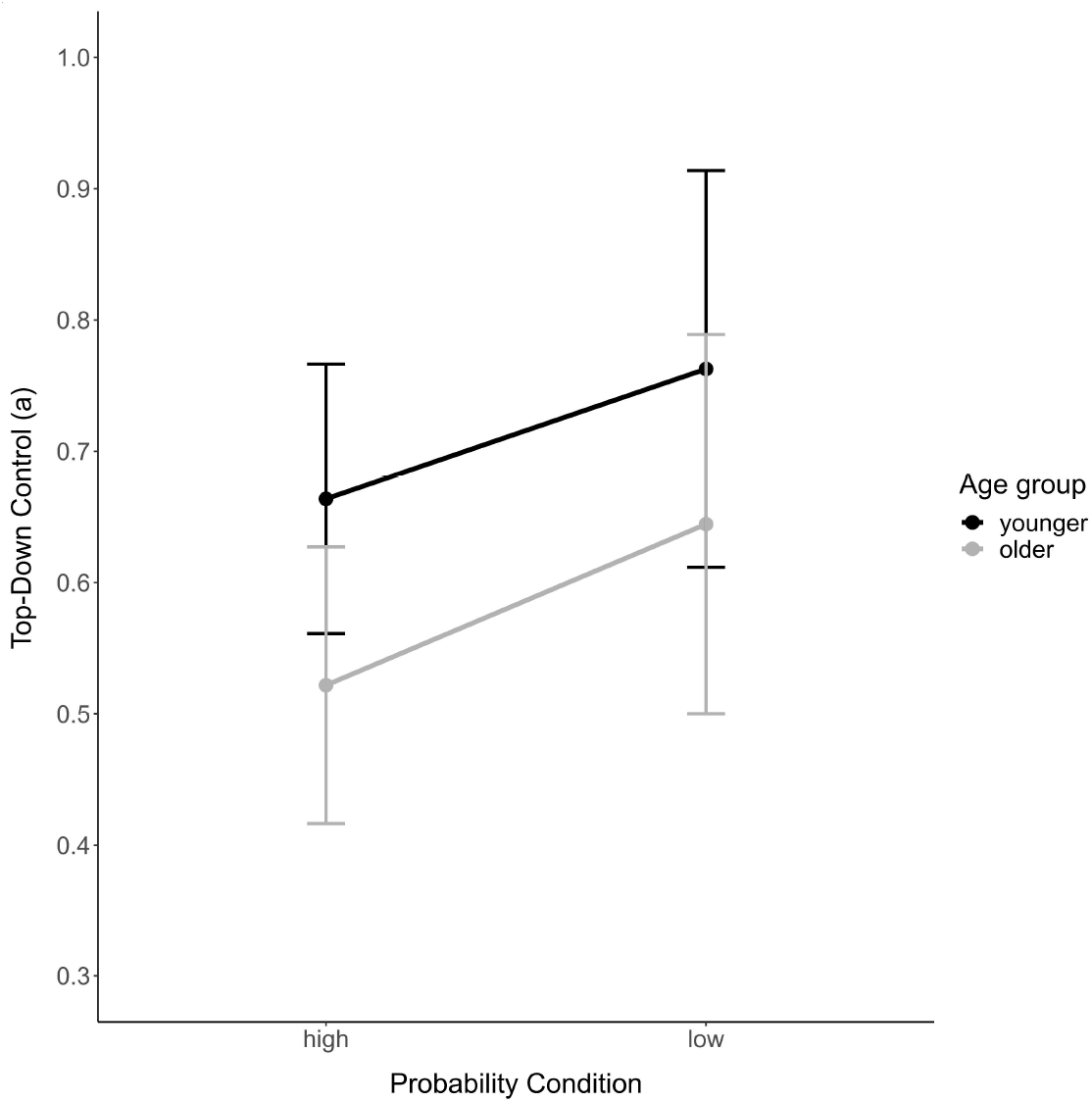
Top-down control in high versus low probability of cue-distractor correspondence in younger and older adults. Note: Only target distractor trials in which the cue indicated the location of an upcoming distractor are included in the analysis. The distractor location was either cued with a high (2/3 or trials) or low (1/3 of trials) probability. Error bars represent confidence intervals.

One fixed stimulus exposure duration, individually determined for each participant, was used for the whole experiment. Exposure durations were adjusted in an initial calibration block and, if necessary, re-adjusted in a subsequent performance check block (see below). While trials of all three display types were shown in the adjustment block, only dual-target trials were used for calibration. Calibration started with an exposure duration of 80 ms, and the adjustment rule was as follows: if the participant was able to report both letters correctly (on dual-target trials), in the next trial the exposure duration was decreased by 10ms; in case no letter was reported correctly, the exposure duration was increased by 10 ms; if only one item could be named, the exposure duration was kept constant. The lowest exposure duration established this way during calibration was then applied to all trials in the subsequent performance check block, which yielded an accuracy estimate for all three types of displays (even though only dual-target trials had been used for the calibration in the preceding block). The aimed-for level of accuracy was 70-90% for single-target trials and above 50% for dual-target trials. If these criteria were not met, the experimenter increased or decreased the exposure duration by 10 ms and re-ran the performance check block. A maximum of three performance check blocks was conducted to determine the final adjusted exposure duration according to the criteria described above.

### Cueing manipulation

The total partial-report task consisted of three blocks. One of these blocks contained 24 single-target and 24 dual-target trials, while the other two blocks contained 48 target-distractor trials each. In the latter (target-distractor) blocks, half of the trials were preceded by an abrupt-onset cue – a 50-ms brightening (i.e., technically: thickening) of the placeholder outline at one location – indicating the upcoming distractor location with 100% validity. Thus, a cue was always followed by a distractor at the same location and never indicated an upcoming target or an empty location. No cue was provided in the other half of trials. Trials with and without cue were presented in randomly intermixed fashion. We hypothesized that abrupt-onset cues would lead to higher (bottom-up) distractor weights and, thus, less efficient estimates of top-down control (parameter α) compared to no-cue trials.

### Procedure

The experiment was conducted in a soundproof and dimly lit cabin. All stimuli were presented on a BenQ 24-inch monitor (1920×1080 pixels screen resolution, 100-Hz refresh rate), at a viewing distance of 60 cm. Head-to-screen distance was controlled by a chin rest, and fixation was monitored by the experimenter using a Logitech c920 HD Pro Webcam mounted on top of the monitor.

The sequence of events on a given trials was as follows: After the initiation of a given trial (manually, by the experimenter), participants were to fixate on the central marker, which was shown together with the four peripheral placeholders for the first 300 ms. Next, the display either remained the same or an abrupt-onset cue was presented at one placeholder location for 50 ms. This was immediately followed by the letter display, which was presented for the individually adjusted exposure duration and then immediately pattern-masked, with the mask being shown for 500 ms. The participant then reported the target letter(s) to the experimenter, who recorded them on the keyboard and then started the next trial with a button press.

At the beginning of the experiment, participants received written instructions on the screen. After reading these and clarifying any issues with the experimenter, they were presented with four example trials (without exposure time limitations) illustrating all possible stimulus conditions and the spatial (distractor) cues. Participants then performed eight training trials at an exposure duration of 120 ms. This was followed by the exposure duration adjustment block and three performance check blocks, to settle on an individual exposure duration which was then introduced in the subsequent three test blocks. Altogether, the experiment lasted around one hour.

### Estimation of distractor and target weights

TVA parameter estimation algorithms employ a maximum-likelihood computation to model partial-report accuracy across all 16 different display configurations (Dyrholm, Kyllingsbæk, Espeseth, & Bundesen, 2011; Kyllingsbaek, 2006). The fitting algorithm yields separate estimates of attentional weights *w_i_* assigned to targets and distractors displayed at each of the four possible locations used in the experiment. Top-down control (parameter α) is defined as the ratio of weights allocated to distractors (*w_D_*) to weights allocated to targets (*w_T_*). Perfect selection without any distractor weighting would result in α equalling 0. In case of non-selective processing, targets and distractors would receive the same weights, yielding an α ratio of 1. Hence, less efficient top-down control (i.e., a strong distractor influence) would be indicated by high α values, whereas efficient top-down control would be indicated by low α values.

### Statistics

We evaluated the data from (preliminary) Experiment 1 using a non-parametric Wilcoxon signed-ranked test.

## Results

Verbal report accuracies for single-target trials, dual-target trials and target-distractor trials with and without preceding cues can be seen in Table 1.

**Table 1.**
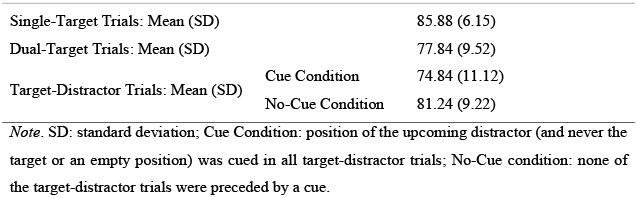
Verbal report accuracy for all participants in preliminary Experiment 1 (N=8)

A Wilcoxon signed-rank test was conducted to analyse the effect of abrupt-onset cues indicating the location of an upcoming distractor. This was done by comparing the top-down control estimates resulting from fitting the data across experimental conditions, once including only target-distractor trials with cued distractors and once including only target-distractor trials without distractor cues. The results revealed a significant difference in top-down control estimates between distractor-cue and no-distractor-cue condition (*Z*= (−2.380), p=.015, r_b_=.944, 95% CIr_b_ [0.762, 0.988]): the distractor weights were higher, i.e. distractors impaired target report performance more, when they where spatially pre-cued (mean α=.861) compared to when they no distractor cue was provided (mean α=.544).

## Discussion

Preliminary Experiment 1 sought to establish that abrupt-onset cues would increase the salience of upcoming distractors at the cued location. As top-down control was less efficient (as indicated by significantly higher α values) when the parameter fitting was performed on the cued target-distractor trials as compared the no-cue trials, the cueing let to the expected higher distractor weights, that is, a competitive advantage of the distractor over the target. The results of this experiment are a prerequisite for Experiment 2, as they show that our cueing manipulation was successful in enhancing the bottom-up salience and, thus, (relative) attentional weight of the (cued) distractor item in a partial-report paradigm.

## Experiment 2

Based on the results of Experiment 1, Experiment 2 was designed to examine whether the (bottom-up) selection advantage conferred to a distractor by the preceding abrupt-onset cue would be top-down modulable by knowledge of the likelihood with which a distractor occurs at the cued location. Would top-down control be improved in a condition in which the distractor appears with a high, as compared to a low, probability at the location of the (salient) abrupt-onset cue? Furthermore, we examined whether the magnitude of any such modulation of top-down control would be influenced (or affected) by healthy ageing. Would the modulation be reduced in older compared to younger adults? In order to address these questions, we varied the probability with which the abrupt-onset cues (the same as used in Experiment 1) indicated the upcoming distractor location in separate trial blocks: blocks with 1/3 of cue-distractor correspondence (i.e., a color-defined distractor appeared with a validity of p = 1/3 at the cued location) versus blocks with 2/3 of correspondence. On trials on which the cue did not indicate the upcoming distractor location, it indicated the target location. Of note, the TVA-based fitting procedure was based exclusively on trials in which the cue indicated the distractor location, permitting us to derive attentional weights for the distractors separately for the high and the low probability condition blocks. Importantly, bottom-up salience of the distractors was equated across these two conditions (as in both conditions the distractor appeared at the cued position). We hypothesized that younger participants would show better shielding against salient distractors when the cue predicted the upcoming distractor location with a high versus a low probability. This would be expressed in better (i.e. lower) estimates of the (top-down control) α parameter the high-probability blocks compared to the low-probability blocks. Furthermore, we wanted to assess whether such active perception effects are modulated in healthy ageing, by comparing older to younger individuals. Arguably, an increased bottom-up attentional focus in ageing should lead to comparable attentional weights for distractors and α (top-down control) parameter estimates in both conditions. If older adults suffer from a decreased ability to use predictive distractor location information, this should result in a significant interaction between group and probability condition.

## Method

### Participants

Thirty-three healthy younger volunteers (≤ 35 years of age) and thirty healthy older volunteers (≥ 60 years) took part in Experiment 2. Three younger and one older participant had to be excluded from analysis because their top-down control values were outliers, differing by > 1.5 times the interquartile range from the upper quartile or the lower quartile (i.e., they were not representative for the respective group). Hence, the final sample consisted of 30 younger and 29 older participants (see Table 2). All participants reported normal or corrected-to-normal vision, including normal colour vision; they all gave informed consent and were either reimbursed for their participation or received course credit. The Edinburgh Handedness Inventory (Olderfield, 1971) was administered to assess handedness. Due to changes in educational and occupational standards over the last decades, we calculated a sociodemographic score (Künstler et al., 2018) based on crystallized intelligence (estimated by a multiple-choice German vocabulary test measuring crystallized intelligence, “Mehrfachwahl-Wortschatz-Intelligenztest”; MWTB; Lehrl, Triebig, & Fischer, 1995), number of school years, and occupation (for a detailed description of the sociodemographic score, see Supplements). Older participants additionally completed the Mini-Mental State Examination (MMSE) as a screening for cognitive impairments indicative of beginning neurodegeneration (Folstein, Folstein, McHugh, & Ingles, 1975). None of the older participants had to be excluded based on a cut-off criterion for cognitive impairment (scores below 27/30 points indicating impairments). Demographic information is presented in Table 2. While both groups differed significantly in age, they were comparable in terms of sex, handedness, and sociodemographic scores. The experiment was approved by the Ethics Committee of the Department of Psychology at LMU München.

**Table 2.** Group demographics in Experiment 2

|  | Younger<br>Participants<br>n =30 | Older Participants<br>n= 29 | Group<br>comparison<br>p value |
| --- | --- | --- | --- |
| Age (years): Mean/SD/Range | 26.6 /4.2/19-34 | 70.8 /4.5/60-77 | <.001 |
| Sex (female/male) | 17/13 | 12/17 | .301 |
| Handedness<br>(right/left/ambidextrous*) | 26/4/0 | 28/0/1 | .353 |
| Sociodemographic Score':<br>Mean/SD/Range | 8.2/1.0/4-9 | 8.0/1.0/6-9 | .438 |
| MMSE score: Mean/SD/Range | / | 28.9/1.1/27-30 | / |

*Note.* SD: standard deviation; handedness: assessed by Edinburgh Handedness Inventory; MMSE: Mini-Mental State Examination, maximum score = 30 points with values < 27 points indicating cognitive impairment; Chi square tests are calculated for handedness and sex, Mann-Whitney U tests are calculated for the other variables;\*due to historical pressure to "relearn" handedness, ambidextrous older participants are treated as left-handed in the statistical comparison.

### Cueing manipulation

The stimuli (including the abrupt-onset cues) and events on a given trial were similar to those in Experiment 1.

One block of trials presented only single- and dual-target displays (randomly intermixed), without any (abrupt-onset) cue. All other blocks exclusively contained target-distractor trials, with the letter display preceded by a cue presented at one of the four possible stimulus locations. On a given trial, the cue appeared either at the location of the subsequent target or the subsequent distractor. Note that the cued location as such was not predictive of where the second stimulus would appear in the display (so the cued location could not be used to infer where the other stimulus would appear). Both trials with and without cue were already introduced during the prior adjustment and performance check blocks, so that participants were accustomed to these types of trial (stimuli) when starting the experiment proper.

The experiment consisted of seven blocks overall. In two high-probability blocks, the abrupt-onset cue indicating the distractor location in two thirds of trials and the target location in the remaining third (overall 144 distractor cue and 72 target cue trials). In contrast, in four low-probability blocks, the distractor location was cued in only one third of the trials and the target in the other two thirds (overall 144 distractor cue and 288 target cue trials). Thus, overall, the two high-probability blocks included the same number of trials on which the cue indicated the upcoming distractor position as the four low-probability blocks. As the subsequent analyses included only target-distractor trials on which the distractor position was cued, allocation of two high- and four low-probability blocks was essential to ensure the same number of trials for statistical comparison. The two high- and the four low-probability blocks were separated by one block of no-cue trials solely presenting single- and dual-target displays (i.e., as there was no cue, a target could never appear at a cued location). The order in which the target-distractor blocks were performed was counterbalanced, with 16 younger and 14 older participants starting with the two high-probability blocks, and the remaining 14 younger and 15 older participants with the low-probability blocks. Participants were not explicitly informed about the probability conditions and the respective variation of likelihood with which a cue indicated the upcoming distractor position; rather, they were to learn the cue-distractor relationship from experience.

### Procedure

The procedure was generally similar to in Experiment 1, except that all target-distractor trials were preceded by a cue, and we introduced a block-wise manipulation of the validity (or probability) with which the cue indicated the distractor location (without telling the participants about this manipulation, so that participants could not know which type of block they were currently performing). The sequence of frames presented on each trial is illustrated in Figure 1A. A short cue-to-letter array SOA (50 ms), brief, individually adjusted exposure durations (younger adults: mean=57.0 ms, SD=16.0 ms, range =30–110 ms; older adults: mean=111.4ms, SD=35.4 ms, range= 60–180 ms) and explicit instructions to maintain fixation over the whole trial episode were intended to prevent any systematic tendency for making eye movements. Additionally, fixation was monitored by camera to ensure that eye movements did not influence performance in a systematic way. Overall, the experiment lasted between one and one and a half hours.

### Estimation of TVA parameters

The estimation of TVA parameters was identical to that described for Experiment 1. TVA parameters were estimated separately for the high-versus low-probability condition. Importantly, only target-distractor trials on which the cue correctly indicated the upcoming distractor location were entered in the fitting procedure (i.e., the fitting was based on the exact same number of such trials in the high- and low-probability cueing conditions). Furthermore, the estimation procedure included the single- and dual-target trials (without any cue) from the intermediate block between the target-distractor blocks for both the high- and the low-probability condition. Thus, separate top-down control parameters α were derived for the two conditions with a high and, respectively, a low probability of the cue correctly indicating the location of an upcoming distractor.

### Statistical analysis

Examining the criterion assumptions for parametric testing in the younger and older participant groups of Experiment 2, a significant Shapiro-Wilk tests indicated a non-normal distribution for top-down control parameter α (W<.880, p<.003) in the high-probability condition for the older group (Shapiro & Wilk, 1965). Accordingly, we calculated a robust repeated-measures ANOVA with the within-participant factor probability (Low vs. High) and the between-participant factor age group (Younger vs. Older) for parameter α. The robust model takes into account that even slight deviations from a normal distribution lead to higher tails of the distribution that substantially increase the standard error of the sample mean. Importantly, there are other location estimators (than means or medians) whose standard errors are significantly less affected by non-normal distributions (Wilcox, 2017). In the present study we chose the sample trimmed mean as location estimator, which is calculated by removing the *x* largest and *x* smallest values in an ascending list of observations depending on the degree of trimming chosen. Generally, a trimming of 20% is recommended (Wilcox, 2017). We followed this recommendation in the present analysis, which was performed using the WRS package (Wilcox & Schönbrodt, 2017) in R Studio version 1.0.136 (RStudio Team, 2016).

In addition, we performed the Bayesian counterpart of repeated-measures ANOVAs (Rouder, Morey, Verhagen, Swagman, & Wagenmakers, 2017) using JASP version 0.8.5 (JASP Team, 2018). To date, a non-parametric, Bayesian counterpart of a repeated-measures design is unavailable and, hence, could not be used. Based on the null and alternative hypotheses, JASP calculates the Bayes factor (B_10_), which is a measure of the ratio of the likelihoods associated with the two hypotheses. Accordingly, if B10 is greater than 3, the data are taken to substantially support the alternative hypothesis, whereas values smaller than 1/3 are interpreted as substantial support for the null hypothesis. B_10_ values between 1 and 3 (as well as between 1 and 1/3 accordingly) would only yield ‘anecdotal evidence’ for an hypothesis (e.g. Dienes, 2011; Wagenmakers et al., 2011).

## Results

Verbal report accuracies for younger and older participants on single-target trials, dual-target trials and target-distractor trials, separately for the high- and low-probability conditions are given in Table 3.

**Table 3.**
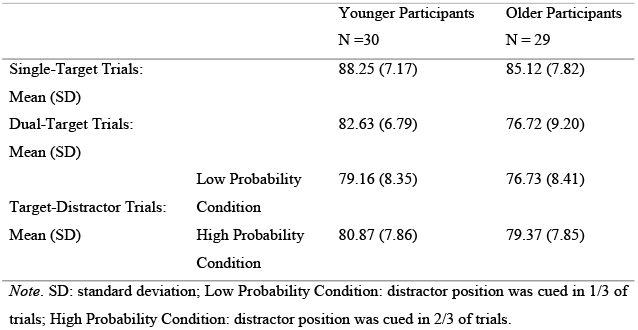
Verbal report accuracy for younger and older participants in Experiment 2

The robust repeated-measures ANOVA on the top-down control parameters α based on trimmed means revealed a significant main effect of probability condition (*Qb*= 6.925, p=.013), but no significant main effect of the between-participant factor age group (Qa= 3.306, p=.078) and no significant interaction (*Qab*= 1.012, p= .322). The main effect of probability condition was due to top-down control being significantly more efficient (as indicated by lower α values) in the condition in which the cue indicated the distractor location with a high probability, as compared to the low-probability condition (.59 vs. .70; see Table 4). The lack of an interaction indicates that the efficiency of top-down control was modulated to a comparable extent by the cue information in both (the younger and older) age groups.

**Table 4.** TVA parameter estimates for top-down control in high versus low probability conditions for younger and older participants

|  |  | Younger Participants<br>N = 30 | Older Participants<br>N = 29 |
| --- | --- | --- | --- |
| Parameter $\alpha$: | Low Probability Condition | 0.76 (0.40) | 0.64 (0.38) |
| Mean (SD) | High Probability Condition | 0.66 (0.27) | 0.52 (0.28) |

*Note.* SD: standard deviation; Low Probability Condition: distractor position was cued in 1/3 of trials; High Probability Condition: distractor position was cued in 2/3 of trials.

The Bayesian repeated-measures ANOVA estimated the observed main effect of probability condition to be 6.039 times more likely under the alternative than the null hypothesis. Furthermore, a Bayes Factor of 0.952 for the age group comparison provided no conclusive support for either (the alternative or the null) hypothesis. And a Bayes Factor of 0.267 for the interaction provided substantial evidence in favour of the null hypothesis (B= 0.267).

## Discussion

Experiment 2 set out to investigate whether younger and older participants could use probabilistic information provided by cues about the (likely) distractor location to shield attentional selection against salient distractors. In order to address this question, we manipulated the probability with which the distractor appeared at the cued location in different trial blocks (1/3 vs. 2/3 of trials). The statistical analyses yielded a significant main effect of probability, indicating more efficient top-down control in blocks in which the cue was highly informative about the upcoming distractor location compared to when its information value was low. This finding suggests that, even though abrupt-onset spatial cues increase the attentional weights of distractors (as demonstrated in Experiment 1), participants can use prior statistical information about the likelihood with which a distractor will appear at an indicated position in order to mitigate this bottom-up effect. Thus, given that the bottom-up-induced salience of the distractors is equal in the two probability conditions, the effective distractor weights are lower in the condition with a high probability of a distractor appearing at the cued location. As the effect was comparable in both age groups, the results suggest that older adults are equally able to utilize predictive cue information in order to minimise attentional capture by salient distractors.

One objection that might be levelled against an interpretation of the main effect of ‘probability’ in terms of a cue-information-induced modulation of the distractor weights (their relative down-modulation in the 2/3 as compared to the 1/3 cue-validity condition) is that this modulation was influenced by the inverse likelihood with which the target appeared at the cued location. Recall that when, in the high-probability condition, the distractor appeared with p = 2/3 at the cued location, the target appeared with p = 1/3 at the cued location, and vice versa in the low-probability condition. However, we contend that this inverse relationship is not a confound, because, in order to perform the task optimally, observers have to set themselves consistently for the target-defining colour (say red, as opposed to the distractor-defining colour blue). Thus, a cue appearing at the target location would render the target more bottom-up salient (whatever the likelihood with which the target appears at the cued location). Critically, however, this would not improve performance differentially between the two cue-validity conditions, because the (positive) top-down set for the target colour is already optimal. By contrast, the task does not necessarily require a (negative) top-down set against the distractor colour, but such a set might be invoked (or be invoked more strongly) when the cue indicates the distractor location with a high, versus a low, probability. In this case, the cue would predict that a distractor-coloured letter would appear at the cued location, so that actively shielding against intrusions of this colour into vSTM would contribute to enhancing target-related selectivity. On this reasoning, one would predict a cue-information-induced modulation of the distractor weights, as evidenced by the results of Experiment 2. But one would not expect a cue-information-induced modulation of the target weights. To examine for the latter, we ran a control analysis including single-target and dual-target trials (both without cues) as well as target-distractor trials in which the cue appeared at the position of the upcoming target (see Supplements). This analysis revealed the estimates of top-down control (based on target-distractor trials in which the upcoming target location was cued) to be comparable between the high- and low-probability blocks (.50 vs. .43 with lower α values indicating better top-down control; see Supplementary Tables S1 and S2), as predicted. This null-effect reinforces the interpretation advanced above, namely, that the probability manipulation in Experiment 2 improved attentional selectivity by lowering the selection weight of the distractor rather than by increasing the weight of the selection weight of the target.

## General Discussion

The present study investigated whether participants can utilise acquired (pre-) knowledge of the location of an upcoming salient distractor to alter their top-down control settings, so as to reduce bottom-up attentional capture. In addition, we examined whether the ability to profit from such predictive information is modulated by healthy ageing. Experiment 1 demonstrated that abrupt-onset spatial cues marking the location of an upcoming distractor increase the distractor’s bottom-up salience and, thus, its weight in the race for being selected by attention. In Experiment 2, we examined whether this bottom-up-induced effect can be actively (down-) modulated, by older as well as younger participants, based on the informativeness of such (abrupt-onset) cues with respect to the location of an upcoming distractor. Comparing top-down control between blocks with high-versus low cue probabilities (the cue predicted the distractor location in 2/3 versus 1/3 of the trials within a block) revealed top-down control to be better in the high-as compared to the low-probability condition. This effect did not differ between younger and older participants: both age groups were similarly able to actively shield attentional selection against bottom-up-induced capture through the (top-down) use of statistical ‘prior’ information.

The results of the present study are in line with previous work suggesting that older participants can utilize predictive information to improve visual search performance (Madden et al., 2014, 2004, Whiting et al., 2007, 2014). Furthermore, they indicate that benefits for top-down control deriving from pre-knowledge are not restricted to probabilistic information regarding the location of an upcoming target, but are also obtained with information about the likely location of an upcoming distractor. The preserved ability of older participants to efficiently use probabilistic information about distractor locations has important implications for seniors’ everyday life: the age-related increased proneness to attentional capture and the attendant costs of distraction can be compensated for better in situations in which the locations of distractors are relatively predictable (as compared to being non-predictable).

Moreover, the present findings are in line with previous reports that younger adults are able to reduce distractor interference and improve top-down control when conditions afford valid predictions about the location of an upcoming distractor (Awh et al., 2003; Chao, 2010; Havlíček et al., 2018; Ruff & Driver, 2006; Sauter et al., 2019, 2018; Wang & Theeuwes, 2018; Watson & Humphreys, 1997). Note, however, that Noonan et al. (2016) have argued that top-down controlled distractor suppression is possible only when observers can make stable predictions, which is not the case with foreknowledge of the distractor location provided by trial-wise presented (i.e., with regard to the indicated location: flexible) cues. According to Noonan et al. (2016), studies that reported behavioural benefits from cues indicating flexibly changing locations of upcoming distractors suffer from various shortcomings in their design, potentially confounding the results obtained. Firstly, unbalanced numbers of trials (Awh et al., 2003) can lead to a higher exposure to distractor-present as compared to distractor-absent trials. In this case, repetition effects could be the underlying cause for more efficient distractor suppression (Noonan et al., 2016). Secondly, binary display configurations with only one possible stimulus location per hemifield (Ruff & Driver, 2006) should be avoided to prevent participants from re-interpreting the distractor location cue as a spatial cue with respect to the location of the upcoming target. Such displays may, conceivably, lead to a re-orientation of attention to the target location, making it impossible to disentangle the underlying – suppression versus facilitation – processes (Noonan et al., 2016). Finally, high similarity of the target and distractor stimuli (Chao, 2010) should be avoided (Noonan et al., 2016). Crucially, the present study design addressed all of the abovementioned points: (i) the exact same number of distractor-cued trials in the low and high probability conditions rules out confounding by potential repetition effects; (ii) in contrast to binary designs, four potential stimulus locations with either horizontal or vertical target-distractor arrangements ensured that uncertainty remained with respect to where the target would appear (in case the distractor was cued); and (iii) the facts that the task did not require participants to make a (speeded) motor response and that the target and distractor letters were defined by a clear feature difference (red vs. blue coloured letters) rule out response criteria and stimulus similarity confounds. Critically, this design – with flexible (trial-by-trial) cues as well as variable distractor locations – still revealed benefits of probabilistic cue information about an upcoming distractor location for the top-down control of (feature-based) stimulus selection.

The present findings have implications for future studies. First, it is quite possible that maintained top-down attentional processes in ageing (as observed in our sample of older participants) are a source of compensation for an underlying stronger susceptibility to bottom-up attentional capture (Madden, 2007; Madden, Spaniol, Bucur, & Whiting, 2007). Neuroimaging findings of age-related increased activations in fronto-parietal structures during the performance of tasks with top-down involvement (Madden, Spaniol, Whiting, et al., 2007) are consistent with such a ‘compensation’ account. Nevertheless, further studies examining top-down control in healthy ageing are needed to uncover the precise underlying mechanisms. Second, a TVA-based study suggests that estimates of top-down control α are decreased in patients with mild cognitive impairment, and deteriorate further in patients with Alzheimer’s disease (Redel et al., 2012). Accordingly, it would be of interest to examine whether the ability to acquire and use statistical pre-knowledge to increase top-down control might be compromised or even absent in patients at risk for pathological ageing and cognitive decline.

## Conclusion

In summary, the present study for the first time adopted a purely perceptual partial-report task based on the TVA and combined it with (abrupt-onset) spatial cues to assess the effect of probabilistic pre-knowledge about the location of an upcoming distractor on top-down control. The results indicate that healthy younger as well as older adults can more effectively shield attentional selection against salient distractors when the probability that a distractor appears at the location of a preceding cue is high as compared to low. This was the case even though the distractors’ bottom-up saliency was enhanced by the cues. This finding suggests that the ability to use the spatial information conveyed by the cues to increase top-down control and, thus, more efficiently shield against attentional capture by a salient distractor is preserved in healthy ageing.

## Acknowled gements

We thank all our participants for taking part in the study.

## Funding

This work was supported by the German Forschungsgemeinschaft (DFG) under Grant FI 1424/2-1 and SO 1336/1-1 as well by the Elite Network of Bavaria under a research scholarship to MH.

## Disclosure Statement

The authors have no actual or potential conflicts of interest.

## Data availability statement

The data are available upon request.

