## SupplementaryMaterial for "Predictability of salient distractor increases top-down control in healthy younger and older adults"

**Sociodemographic Score**

See:	Künstler, E.C.S., Penning, M.D., Napiórkowski, N., Klingner, C.M., Witte, O.W., Müller, H.J., Bublak, P., Finke, K., 2018. Dual task effects on visual attention capacity in normal aging. Front. Psychol. 9, 1–12. https://doi.org/10.3389/fpsyg.2018.01564

Due to changes in educational and occupational standards over the past decades, we created a sociodemographic score based on an estimate of crystallized intelligence, number of school years, and occupation (either intended or obtained). A maximum of 3 points could be obtained per category, yielding an overall maximum score of 9 points.

Crystallized intelligence was estimated by administering the “Mehrfachwahl-Wortschatz-Intelligenztest”; MWTB; Lehrl, Triebig, & Fischer, 1995), a multiple-choice German vocabulary test. The points were allocated as follows: 1 point for those below average (estimated IQ < 85) ; 2 points for those with an average score (estimated IQ = 100 ± 15); and 3 points for those with an above average score (estimated IQ > 115) according to the MWTB handbook.

Secondary school qualifications were categorized as follows: Participants completing a qualification which required 9 years of schooling obtained 1 point; those who completed a qualification which necessitates 10 years of education were awarded 2 points; and those who had a qualification which required 12 school years were given 3 points.

Finally, participants were scored according to their occupation: 1 point was given to those participants with menial jobs which did not require any further training or education; 2 points were given to those whose occupation required further training; 3 points were awarded to those participants with occupations requiring a university degree. University students were automatically awarded 3 points, even if they had not as yet completed their degree.

**Explorative analysis: Target-Distractor trials in which the upcoming target location was cued**

The explorative analysis (including only target-distractor trials in which the cue indicated the upcoming target location) yielded no significant main effect of probability condition (high vs. low), main effect of age group (younger vs. older) or interaction effect (all F≤ 2.977, all p ≥ .090). The comparison of probability conditions substantially favours the null hypothesis B_10_= 0.214). A Bayes Factor of B_10_= 0.333 for the age group comparison provides anecdotal evidence for equal group means. The interaction effect yields an inconclusive Bayes Factor of B_10_= 0.949. The results suggest that estimates of top-down control are comparable over target-cue trials, not significantly differing between age groups or based on the probability condition.^[1]^

| Table S1.  *Verbal report accuracy for younger and older participants when only including target-distractor trials in which the abrupt-onset cue indicated the upcoming target location.* | | | |
| --- | --- | --- | --- |
|  |  | Younger Participants | Older Participants |
|  |  | N =30 | N = 29 |
| Target Distractor Trials: Mean (SD) | Low Probability Condition | 85.28 (7.77) | 78.06 (10.98) |
|  | High Probability Condition | 86.77 (5.85) | 77.30 (10.60) |
| *Note*. SD: standard deviation; Low Probability Condition: target position was cued in 1/3 of trials; High Probability Condition: target position was cued in 2/3 of trials. | | | |

| Table S2.  *TVA parameter estimates for younger and older participants based on target-distractor trials in which the cue indicated the upcoming target position* | | | |
| --- | --- | --- | --- |
|  |  | Younger Participants | Older Participants |
|  |  | N =30 | N = 29 |
| Parameter α:  Mean (SD) | Low Probability Condition | 0.42 (0.25) | 0.44 (0.22) |
|  | High Probability Condition | 0.48 (0.33) | 0.52 (0.30) |
| *Note*. SD: standard deviation; 3 older participants had to be excluded because their top-down control values were outliers that were not representative for the respective group (differing by > 1.5 times the interquartile range from the upper quartile or the lower quartile); Low Probability Condition: target position was cued in 1/3 of trials; High Probability Condition: target position was cued in 2/3 of trials. | | | |

1. We acknowledge that this null-effect should be interpreted with caution as only 72 target-distractor trials with target cues could be fed into the fitting algorithm per condition, rendering the resulting estimates of top-down control parameter α somewhat less reliable compared to the inverse conditions in which the distractor location was cued.
